# Genetic and epigenetic insight into morphospecies in a reef coral

**DOI:** 10.1101/119156

**Authors:** James L. Dimond, Sanoosh K. Gamblewood, Steven B. Roberts

## Abstract

Incongruence between conventional and molecular systematics has left the delineation of many species unresolved. Reef-building corals are no exception, with phenotypic plasticity among the most plausible explanations for alternative morphospecies. As potential molecular signatures of phenotypic plasticity, epigenetic processes may contribute to our understanding of morphospecies. We compared genetic and epigenetic variation in Caribbean branching *Porites* spp., testing the hypothesis that epigenetics— specifically, differential patterns of DNA methylation—play a role in alternative morphotypes of a group whose taxonomic status has been questioned. We used reduced representation genome sequencing to analyze over 1,000 single nucleotide polymorphisms and CpG sites in 27 *Porites* spp. exhibiting a range of morphotypes from a variety of habitats in Belize. We found stronger evidence for genetic rather than epigenetic structuring, identifying three well-defined genetic groups. One of these groups exhibited significantly thicker branches, and branch thickness was a better predictor of genetic groups than depth, habitat, or symbiont type. Epigenetic patterns were more subtle, with no clear groups. The more thickly branched individuals in one of the genetic groups exhibited some epigenetic similarity, suggesting potential covariation of genetics and epigenetics. This covariation was further supported by a positive association between pairwise genetic and epigenetic distance. We speculate that epigenetic patterns are a complex mosaic reflecting inheritance and diverse environmental histories. Given the role of genetics in branching *Porites* spp. morphospecies we were able to detect with genome-wide sequencing, use of such techniques throughout the geographic range may help settle their phylogeny.

## Introduction

Inconsistencies between conventional and genetics-based taxonomy are common across taxa (Patterson *et al.* 1993). Morphological characters have long been the basis of systematics, yet they can be misleading in cases such as convergent evolution or extensive phenotypic plasticity (Potter *et al.* 1997; Fukami *et al.* 2004; Fritz *et al.* 2007). While molecular phylogenetics has illuminated species relationships by identifying and resolving some of these issues, uncertainties often persist, and disagreement among molecular studies is not uncommon. The inferences made by molecular studies are influenced by numerous factors such as the number and type of markers and the models used to analyze them (Brocchieri 2001; Yang & Rannala 2012). Moreover, the field of epigenetics has uncovered novel mechanisms of phenotypic variation and inheritance that has led to reappraisal of the traditional theory of molecular evolution (Noble 2015; Skinner 2015).

Reef-building corals exhibit substantial inter- and intraspecific variation in growth forms that has proved particularly problematic for the taxonomic delineation of species (Fukami *et al.* 2004; Forsman *et al.* 2009; Flot *et al.* 2011). Conventional taxonomy based on morphological features has obscured coral phylogenies and in some cases overestimated species diversity revealed by way of genetic analyses (Fukami *et al.* 2004; Forsman *et al.* 2010; Prada *et al.* 2014). In other cases, supposed ecomorphs of the same species have turned out to be sibling species (Knowlton *et al.* 1992), and genetic data have also uncovered cryptic species among similar growth forms of what was thought to be a single species (Keshavmurthy *et al.* 2013; Schmidt-Roach *et al.* 2013). The confusion surrounding coral taxonomy has consequences for coral conservation, because a species cannot be conserved and managed if it cannot be defined. For example, 25 coral species are currently listed under the U.S. Endangered Species Act (NOAA 2015), and the ambiguous taxonomic status of some of these species has necessitated revision using molecular data (Forsman *et al.* 2010).

Several studies have concluded that morphological plasticity is among the most plausible hypotheses for instances of both over- and underestimation of species diversity based on conventional taxonomy (Forsman *et al.* 2010; Flot *et al.* 2011; Keshavmurthy *et al.* 2013; Prada *et al.* 2014). Indeed, a recent reciprocal transplant study confirmed the role of phenotypic plasticity in alternative morphotypes of *Pocillopora* spp. (Paz-García *et al.* 2015). Morphological plasticity is likely to be particularly prevalent and adaptive in corals due to their sessility, longevity, modularity, and indeterminate growth (Jackson & Coates 1986; Sebens 1987; Todd 2008). In a review of 26 studies that tested for morphological plasticity in approximately 20 different species of reef corals, 92% of studies documented evidence for plasticity (Todd 2008).

The expanding field of epigenetics may hold promise for understanding phenotypic plasticity in corals. Epigenetic processes are increasingly recognized as molecular signatures of phenotypic variation (Duncan *et al.* 2014). DNA methylation, the most widely studied epigenetic mark, involves the addition of a methyl group to a cytosine, most commonly in the context of a CpG dinucleotide pair. Along with other epigenetic features such as histone modifications and small RNA molecules, methylation patterns can influence gene expression, though the mechanisms appear to be diverse, complex, and context-dependent (Gavery & Roberts 2014; Duncan *et al.* 2014). Unlike the DNA sequences of the genome itself, which change relatively little during an individual’s lifetime, genome DNA methylation patterns are not fixed, and can be influenced by environmental stimuli (Duncan *et al.* 2014). An expanding number of studies have documented differential patterns of DNA methylation associated with alternative phenotypes in a range of taxa (Kucharski *et al.* 2008; Fonseca Lira-Medeiros *et al.* 2010; Smith *et al.* 2015, 2016; Schield *et al.* 2016). Among different morphotypes of mangroves living in distinct habitats, for example, (Fonseca Lira-Medeiros *et al.* 2010) found little genetic variation but high levels of epigenetic variation. In the genomes of two alternative morphotypes of threespine sticklebacks, (Smith *et al.* 2015) identified 77 differentially methylated regions whose functions were associated with known adaptive phenotypes. Interestingly, even among different species of Darwin’s finches, epigenetic differences were better correlated with traditional phylogenetic relationships than were genetic differences (Skinner *et al.* 2014). Thus, an increased understanding of epigenetics may revise definitions of biological diversity and evolution (Skinner 2015), with implications for future conservation efforts.

The taxonomic status of branching corals of the genus *Porites* in the tropical Western Atlantic has been uncertain. There are currently three recognized species: *P. porites* (Pallas, 1766), *P. furcata* (Lamarck, 1816), and *P. divaricata* (Lesueur, 1820). Branch diameter and corallite features have traditionally been used to define these species, and while one study found morphological variation to be nearly continuous (Brakel 1977), other studies have identified morphological breaks, with some supporting traditional taxonomic delineations (Weil 1992; Budd *et al.* 1994; Jameson 1997; Jameson & Cairns 2012). Molecular studies have also failed to reach consensus (Weil 1992; Budd *et al.* 1994; Forsman *et al.* 2009; Prada *et al.* 2014), but earlier studies used allozymes, which could be influenced by epigenetic effects. The most recent and thorough population genetic analysis found no support for upholding the three named species, finding no significant variation between supposed species across 11 genetic markers and multiple geographic sub-regions (Prada *et al.* 2014). The authors acknowledged that it is possible that their study overlooked diversity somewhere in the genome, but identified phenotypic plasticity as a plausible explanation for their results. To further explore the potential source of phenotypic variation in branching *Porites* spp., we compared genetic and epigenetic diversity in these corals using restriction site associated DNA (RAD) sequencing.

## Methods

### Specimen collection and DNA extraction

Corals were collected in May 2016 within approximately 4 km of the Smithsonian Institution’s Carrie Bow Cay Field Station in Belize (16° 48’ 9.39” N, 88° 4’ 54.99” W). In the field, the three Caribbean branching *Porites* spp. are distinguished primarily by branch diameter and habitat type, so collections targeted a broad range of branch diameters (6 - 26 mm), depths (0.5 - 17 m) and habitats (mangrove to forereef) to sample as much variation as possible. A total of thirty specimens were collected, but we obtained sufficient sequence data from only 27 of these (Table 1). Branch tips were cut with shears and placed in a conical tube. Within one hour of collection, specimens were transferred to tubes containing salt-saturated DMSO (SS-DMSO) for DNA preservation (Gaither *et al.* 2011). Coral tissue preserved in SS-DMSO began to slough off the skeleton after several days and was easily removed using forceps. Small pieces of tissue (∼0.5 μl volume) were washed three times via centrifugation with phosphate buffered saline prior to DNA extraction using Qiagen DNeasy Blood and Tissue kits according to the manufacturer’s protocol, with an overnight lysis with proteinase K. Samples were further purified via overnight ethanol precipitation, followed by resuspension in Qiagen AE buffer. Genomic DNA was checked for yield and quality via fluorescence (Qubit BR assay) and gel electrophoresis, respectively.

**Table 1.** *Porites* spp. specimen data. For descriptions of habitat types on the Belize Barrier Reef refer to Rützler & Macintyre (1982).

| Sample # | Coordinates | Habitat | Depth (m) | Symbiont clade | Branch diameter (mm) |
| --- | --- | --- | --- | --- | --- |
| 101 | N 16° 48.126', W 88° 4.729' | Lower spur & groove | 7.9 | C | 19.5 |
| 103 | N 16° 48.294', W 88° 4.666' | Inner reef slope | 16.2 | C | 10.5 |
| 104 | N 16° 48.294', W 88° 4.666' | Inner reef slope | 16.5 | C | 9 |
| 105 | N 16° 46.792', W 88° 4.599' | Lower spur & groove | 9.8 | C | 11.5 |
| 106 | N 16° 46.792', W 88° 4.599' | Lower spur & groove | 7.0 | C | 17 |
| 107 | N 16° 46.821', W 88° 4.521' | Inner reef slope | 12.5 | C | 11.5 |
| 108 | N 16° 48.120', W 88° 4.751' | Upper spur and groove | 7.0 | NA | 10.5 |
| 109 | N 16° 46.821', W 88° 4.521' | Inner reef slope | 14.6 | C | 10 |
| 110 | N 16° 48.146', W 88° 4.658' | Inner reef slope | 14.0 | C | 14 |
| 111 | N 16° 48.120', W 88° 4.751' | Upper spur and groove | 7.9 | C | 14 |
| 112 | N 16° 46.571', W 88° 4.524' | Lower spur & groove | 8.5 | C | 13 |
| 114 | N 16° 46.798', W 88° 4.511' | Outer ridge | 15.5 | C | 9 |
| 115 | N 16° 48.354', W 88° 4.998' | Backreef | 1.2 | C | 20 |
| 116 | N 16° 48.354', W 88° 4.998' | Backreef | 1.2 | NA | 10 |
| 117 | N 16° 48.354', W 88° 4.998' | Backreef | 1.2 | A | 13 |
| 118 | N 16° 49.645', W 88° 6.364' | Mangrove | 0.6 | C | 7 |
| 121 | N 16° 49.645', W 88° 6.364' | Mangrove | 0.6 | C | 7.5 |
| 122 | N 16° 49.645', W 88° 6.364' | Mangrove | 0.6 | C | 6 |
| 123 | N 16° 48.181', W 88° 4.885' | Backreef | 0.8 | C | 21 |
| 124 | N 16° 48.115', W 88° 4.983' | Lagoon/cut | 4.6 | C | 16.5 |
| 125 | N 16° 46.798', W 88° 4.511' | Outer ridge | 16.5 | C | 10 |
| 126 | N 16° 46.798', W 88° 4.511' | Outer ridge | 17.4 | C | 26 |
| 127 | N 16° 48.115', W 88° 4.983' | Lagoon/cut | 2.7 | A | 15 |
| 128 | N 16° 48.181', W 88° 4.885' | Backreef | 0.6 | NA | 10 |
| 129 | N 16° 48.115', W 88° 4.983' | Turtlegrass | 0.9 | A | 10.5 |
| 130 | N 16° 48.181', W 88° 4.885' | Backreef | 0.8 | C | 17.5 |
| 131 | N 16° 48.181', W 88° 4.885' | Turtlegrass | 0.5 | A | 10 |

The remaining coral skeletons were soaked in a 10% bleach solution for 24 hrs, then dried at 40°C for 48 hrs. The diameter of branch tips at their widest cross-section was measured to the nearest 0.5 mm with vernier calipers.

### Symbiont genotyping

Reef corals commonly engage in species-specific associations with symbiotic dinoflagellates (*Symbiodinium* spp.), and these associations can also be related to genetic structure within a given host (Bongaerts *et al.* 2010; Finney *et al.* 2010). To identify the dominant *Symbiodinium* type associated with each coral, an approximately 700 base pair region of domain V of the cp23S-rDNA region was PCR-amplified using primer pair 23S1 (5’-CACGACGTTGTAAAACGACGGC TGTAACTATAACGGTCC-3’) and 23S2 (5’-GGATAACAATTTCACACAGGCCATCGTATTGAACCCAGC-3’) (Santos *et al.* 2002). PCR was performed in 25 μl volumes containing 1X green buffer (Promega), 2.5 mM MgCl_2_, 240 μΜ dNTP, 5 pmol of each primer, 1U *Taq* (GoTaq, Promega), and 1-20 ng of template DNA. Reactions were carried out in an Applied Biosystems Veriti thermocycler under the following conditions: initial denaturing period of 1 min at 95 °C, 35 cycles of 95 °C for 45 s, 55 °C for 45 s, and 72 °C for 1 min, and a final extension period of 7 min. PCR products were cleaned (NEB Monarch kit) and checked on a 1% agarose gel. Cleaned PCR products (10 ng/μL) were sent to Sequetech Corporation (Mountain View, CA) for Sanger sequencing using the forward primer. Chromatograms were edited with Geneious v9.1.5, converting bases with a Phred score < 30 to Ns. Sequences were then aligned using ClustalW and queried against the GenBank nucleotide database in Geneious.

### RAD library preparation and sequencing

Double digest RADseq (ddRADseq) libraries were prepared following the methods of (Peterson *et al.* 2012). Only samples with high molecular weight DNA were used for library preparation. For each sample, a minor variation of ddRADseq called EpiRADseq was also employed to evaluate methylated loci (Schield *et al.* 2016). Both methods use two restriction enzymes—a rare cutter and a common cutter—to perform a double digest of the DNA at specific restriction sites throughout the genome, with the only difference that EpiRADseq uses a methylation-sensitive common cutter that will not cut methylated loci. Both methods used the rare cutter *PstI* (5’-CTGCAG-3’ recognition site), while the common cutter *MspI* (5’-CCGG-3’ recognition site) was used for ddRADseq and the methylation-sensitive isoschizomer *Hpaii* (also 5’-CCGG-3’ recognition site) was used for EpiRADseq preparations. Double digests of 300-500 ng gDNA per sample were carried out using 20 units of each enzyme in the manufacturer’s supplied buffer (New England Biolabs, NEB) for 5 hours at 37 °C. Samples were cleaned using magnetic beads (Sera-Mag SpeedBeads) prior to ligation of barcoded Illumina adapters onto the fragments (Peterson *et al.* 2012). After two rounds of bead cleanup, samples were pooled into 12 libraries (along with 18 other samples not analyzed in this study), which were then subjected to automated size-selection of fragments between 415 and 515 bp using a Pippin Prep (Sage Science). Libraries were then PCR amplified using Phusion *Taq* (NEB) and Illumina-indexed primers (Peterson *et al.* 2012). Final library fragment sizes and concentrations were evaluated with D1000 ScreenTape on an Agilent 2200 TapeStation. Libraries were sent to the Vincent J. Coates Genomics Sequencing Laboratory at the University of California, Berkeley, where their concentrations were verified via qPCR prior to 100 bp, paired-end sequencing in equimolar ratios on the Illumina HiSeq 4000.

### RAD sequence assembly

Sequences were assembled using *ipyrad* v0.3.41 (Eaton 2014). We used the ‘denovo - reference’ assembly method with the *Symbiodinium minutum* (clade B; GenBank accession GCA_000507305.1 (Shoguchi *et al.* 2013)) and *Symbiodinium kawagutii* (clade F; http://web.malab.cn/symka_new/data/Symbiodinium_kawagutii.assembly.935Mb.fa.gzb.fa.gz (Lin *et al.* 2015)) genomes used as reference to subtract symbiont reads from the *de novo* assembly. We concatenated these genomes into a single reference file. Step one of the *ipyrad* workflow demultiplexed the data in each pool by identifying restriction overhangs and barcode sequences associated with each sample; zero barcode mismatches were tolerated. Demultiplexed samples were then combined in a single directory for further steps. In step two, reads were trimmed of barcodes and adapters and quality filtered using a q-score threshold of 20, with bases below this score converted to Ns and any reads with more than 5 Ns removed. Step three mapped reads to the concatenated symbiont reference genomes with *BWA* using the default *bwa mem* setting and removed any mapped reads. With the remaining reads, similar clusters of reads were identified using a threshold of 85% similarity and aligning them. We chose 85% as a moderately conservative clustering threshold to avoid over-splitting of loci (Harvey *et al.* 2015). Next, step four performed joint estimation of heterozygosity and error rate (Lynch 2008) based on a diploid model assuming a maximum of 2 consensus alleles per individual. Step five used the parameters from step four to determine consensus bases calls for each allele, and removed consensus sequences with greater than 5 Ns per end of paired-end reads. With consensus sequences identified, step six clustered and aligned reads for each sample to consensus sequences. Finally, step seven filtered the dataset according to maximum number of indels allowed per read end (8), maximum number of SNPs per locus (20), maximum proportion of shared heterozygous sites per locus (0.5), and minimum number of samples per locus (15).

Henceforth, we will use the term *locus* to refer to a consensus paired-end read. The term *SNP* refers specifically to a single nucleotide polymorphism on a locus, while the term *CpG* refers to a cytosine-guanine dinucleotide pair that can be either methylated or non-methylated at the 5’-CCGG-3’ restriction site of each locus.

### SNP analysis

We analyzed unlinked SNPs that were sampled by *ipyrad* at 1 SNP per locus with the least amount of missing data; SNPs were sampled randomly if they had equal amounts of missing data. We were able to estimate the SNP error rate, defined as the proportion of SNP mismatches between pairs of the same individuals (Mastretta-Yanes *et al.* 2015), by treating ddRADseq and EpiRADseq samples as technical replicates and computing pairwise differences between individuals using the *dist.gene* function in the R package *ape* (Paradis et al. 2004). We then analyzed a SNP dataset from ddRADseq libraries that had no missing data across samples. While this reduced the dataset considerably, it allowed us to test variation across a similar set of loci for both the genetic and epigenetic analysis, as explained further below.

Genetic differentiation among samples was examined using multidimensional scaling (k = 2) using the *cmdscale* R function. Discriminant analysis of principal components (DAPC) in the R package *adegenet* (Jombart 2008) was used to further examine these patterns. Rather than applying prior assumptions about the identity of each coral specimen to assign population groups, we used the *find.clusters* function in *adegenet* to identify groups. This k-means clustering function reduces the data with principal components analysis (PCA) before estimating the number of clusters with the lowest Bayesian information criterion (BIC). All PCs were retained for the analysis and a maximum of 10 clusters was specified. Once groups were identified, cross-validation (*xvalDapc*) was used to estimate the number of PCs to retain for the subsequent DAPC analysis. In the groups identified by the DAPC analysis, genetic differentiation between groups was evaluated by computing pairwise Weir and Cockerham’s *F_ST_* using the R package *hierfstat* (Goudet 2005).

### DNA methylation analysis

Methylation detection with EpiRADseq data involves analysis of read counts (Schield *et al.* 2016). Presence of a given read in the EpiRADseq dataset indicates that the locus is not methylated at the restriction cut site. Conversely, if a locus is methylated at the 5’-CCGG-3’ cut site, *Hpaii* is blocked from cutting, and the locus will be absent from the dataset. Hence, read counts of zero are informative, but they could also be the result of missing data through processes such as allele dropout or variation in library size or fragment size selection. To control for this, we used ddRADseq data to normalize the EpiRADseq data. Any loci with zeros in the ddRADseq library were treated as absent and removed, thereby leaving zeros in the EpiRADseq library only where the locus was counted in the ddRADseq library. This had the added benefit that it resulted in analysis of a similar set of loci as the SNP analysis described above.

EpiRADseq and ddRADseq read counts per locus were highly correlated, with the exception of methylated loci, which, as expected, had low or zero abundances in EpiRADseq libraries (Fig. 1A). The dataset comprising read counts for both EpiRADseq and ddRADseq was subject to normalization of read counts using TMM normalization in the R package *edgeR* (Robinson *et al.* 2010). Residuals of linear regressions of EpiRADseq and ddRADseq libraries provided optimal differentiation of methylated loci from non-methylated loci (Fig. 1B&C). Finally, a binary dataset was created from these data by setting a residual threshold of −1 for methylated loci such that 1 = methylated and 0 = non-methylated (Fig. 1C). These data were evaluated using multidimensional scaling as described above for the SNP analysis.

**Fig. 1.**
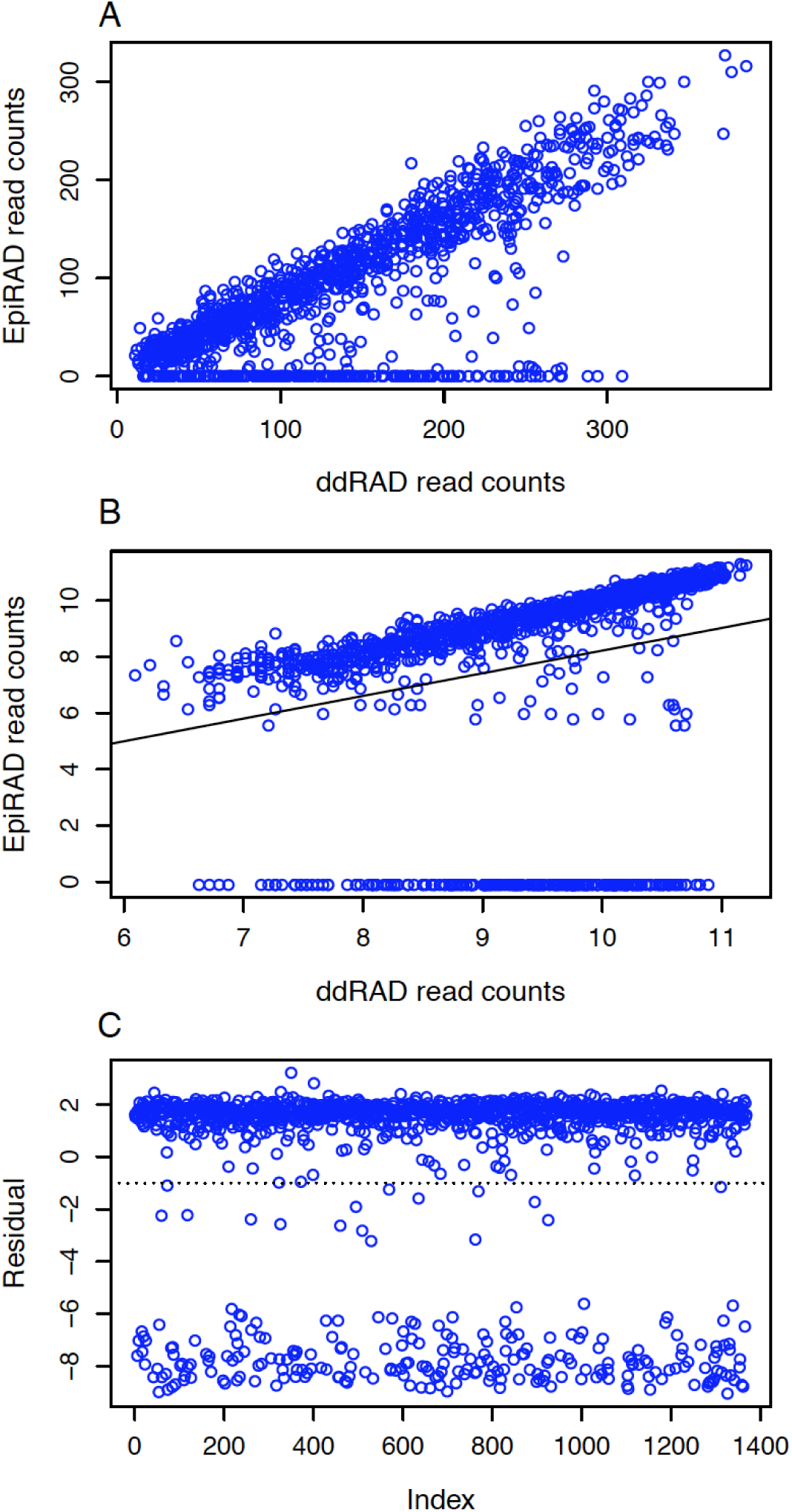
Determining methylated loci in a representative sample. (A) ddRADseq and EpiRADseq raw read counts for each locus (each point is a locus) were highly correlated except for methylated reads, which were absent or at low abundance in the EpiRAD library and cluster at the bottom of the y-axis. (B) ddRADseq and EpiRADseq read counts after TMM normalization, showing regression line from linear model used for derivation of residuals. (C) Residuals from the linear regression of the normalized data. Non-methylated reads have positive residuals and cluster at the top while methylated reads have negative residuals and cluster at the bottom. The binary dataset derived from these data was based on a residual threshold of ⍰ −1 (dotted line) for designating loci as methylated.

EpiRADseq read counts could theoretically represent variable levels of methylation within a sample, such as through variation in methylation across pooled replicates, or among different tissue or cell types (Schield *et al.* 2016). However, our analysis of ddRADseq and EpiRADseq read counts in tandem indicated that much of the variability in read counts occurs in both libraries and is thus likely related to library preparation effects such as fragment size-selection or PCR bias (Davey *et al.* 2013). In the original EpiRADseq method developed by Schield *et al.* (2016), PCR effects were minimized by using unique molecular identifier sequences attached to fragments. We suspect that fragment size-selection effects could be equally important in driving read count variability. For example, abundant reads could simply represent the mean or mode fragment size within the library after size-selection, while less abundant reads could represent the tails of the distribution. In either case, our analysis suggests that much of the read count variation above zero does not reflect methylation levels, and by normalizing EpiRADseq data to the ddRADseq data and creating a binary dataset, we removed much of the potential bias described above.

A repository with the complete bioinformatic workflow described above can be accessed at https://github.com/jldimond/Branching-Porites.

## Results

### Data yield

An average of 3.24 million paired-end reads per sample were obtained based on restriction overhangs and barcodes, with an average of 2.63 million reads per sample remaining after q-score and adapter filtering. After filtering for symbionts and minimum read depth, an average of 26,330 consensus loci per sample were obtained, with a total of 135,980 unique consensus loci across samples. Obtaining datasets with no missing data across samples substantially reduced the number of loci; the final SNP dataset consisted of 1,113 unlinked SNPs (1,113 consensus paired-end loci with 1 SNP sampled per locus) across 27 samples, while the methylation dataset consisted of 1,712 CpGs (1,712 consensus paired-end loci with one CpG cut site per locus) across 25 samples. Differences in the number of SNPs/CpGs between datasets were due to the different numbers of samples; two samples, 101 and 112, were removed from the methylation dataset due to low coverage. This reduced the number of samples in the methylation dataset but resulted in a greater number of shared CpG sites.

### Symbiont identity

BLAST searches of *Symbiodinium* cp23S sequences indicated that the dominant symbiont in the majority of corals was a clade C *Symbiodinium* with high similarity to *Symbiodinium* subclade C3 (Table 1). Four corals occurring in shallow water habitats (⍰ 2.7 m) hosted clade A *Symbiodinium* with high similarity to subclade A3. We were unable to amplify cp23S in three of the specimens despite repeated attempts.

### Genetic patterns

Based on SNP mismatches between technical replicates, the SNP error rate was estimated to be 3.6% (standard deviation, SD, 3.1%), which is on the lower end of the range that has been reported previously (Mastretta-Yanes *et al.* 2015) (Fig. 2). In other words, multilocus genotypes of technical replicates were 96.4% similar on average. In contrast, nearly all non-replicate pairwise comparisons exhibited greater differences, with the exception of two pairs of individuals (109 & 114, 127 & 115) that exhibited similarity within the range of the SNP error (Fig. 2). The high similarity of these individuals suggests they are possible clones. Specimens 109 and 114 were separated by approximately 45 m, while specimens 127 and 115 were separated by approximately 450 m. If they are clones, these distances make it unlikely that they resulted from fragmentation.

**Fig. 2.**
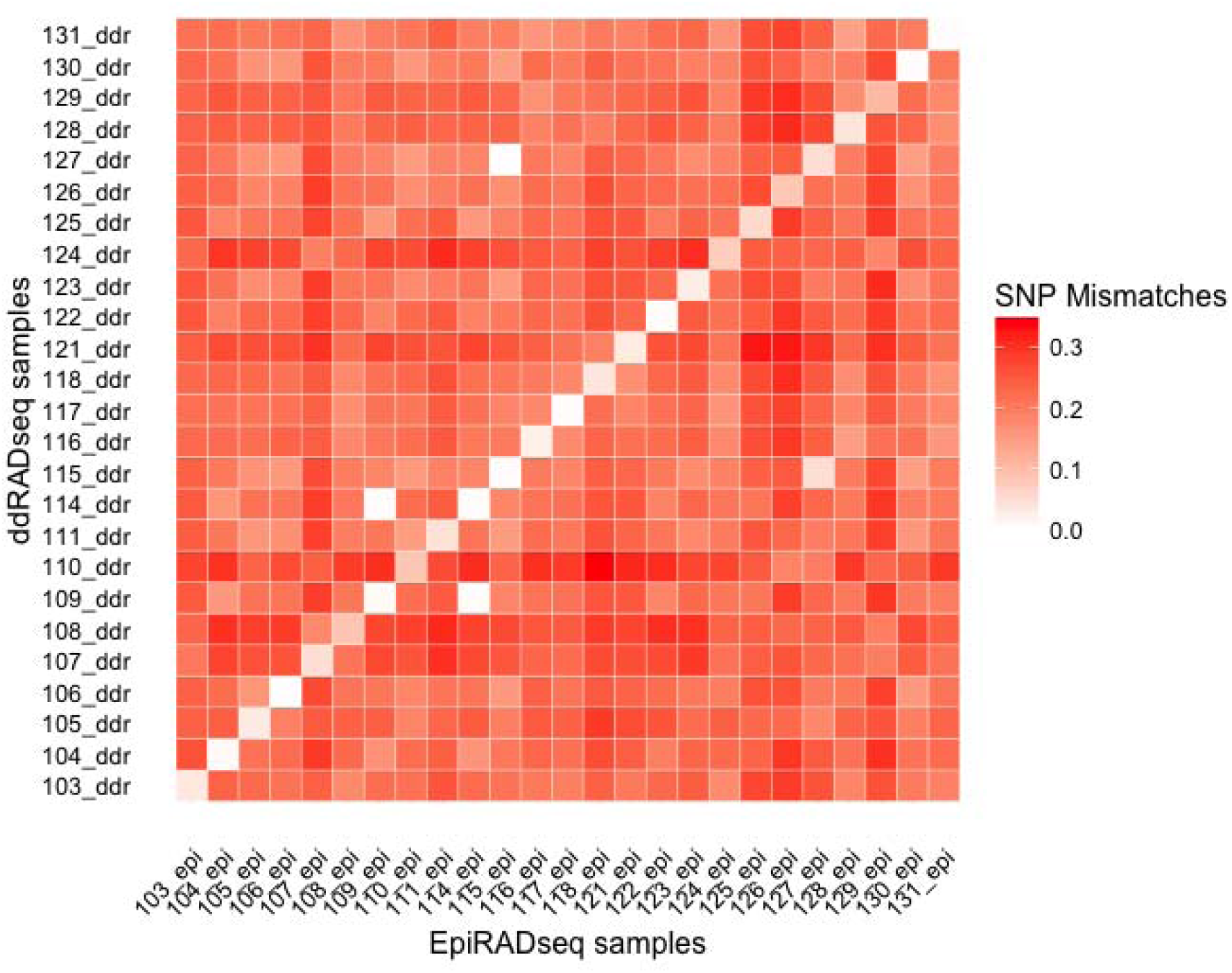
Pairwise comparisons of SNP mismatches between all 25 individuals with both ddRADseq and EpiRADseq data. The color ramp represents the proportion of SNP mismatches, also known as genetic distance, between pairs. ddRADseq and EpiRADseq data for the same individual were used as technical replicates to estimate the SNP error rate and are expressed along the diagonal. Two pairs of outliers with a low proportion of mismatches (109/104, 115/127) are possible clones.

Multidimensional scaling of samples based on SNPs suggested genetic structure of *Porites* spp., with samples clustering into three generally well-separated groups (Fig. 3). K-means clustering of SNPs showed the strongest support for three clusters according to BIC values (Fig. 4A). Based on results of cross-validation, nine PCs (the maximum suggested given the sample size) were retained for DAPC analysis of the three groups, which resulted in two discriminant axes that were both retained (Fig. 4B inset). As with the MDS analysis, DAPC indicated clear separation of the three SNP groups (Fig. 4B). The two pairs of potential clones were in two separate groups (109 & 114 in group 2, 127 & 115 in group 1). Similar values of *F_ST_* were observed between groups (*F_ST_* for each pairwise comparison: 1&2 = 0.194; 1&3 = 0.207; 2&3 = 0.191).

**Fig 3.**
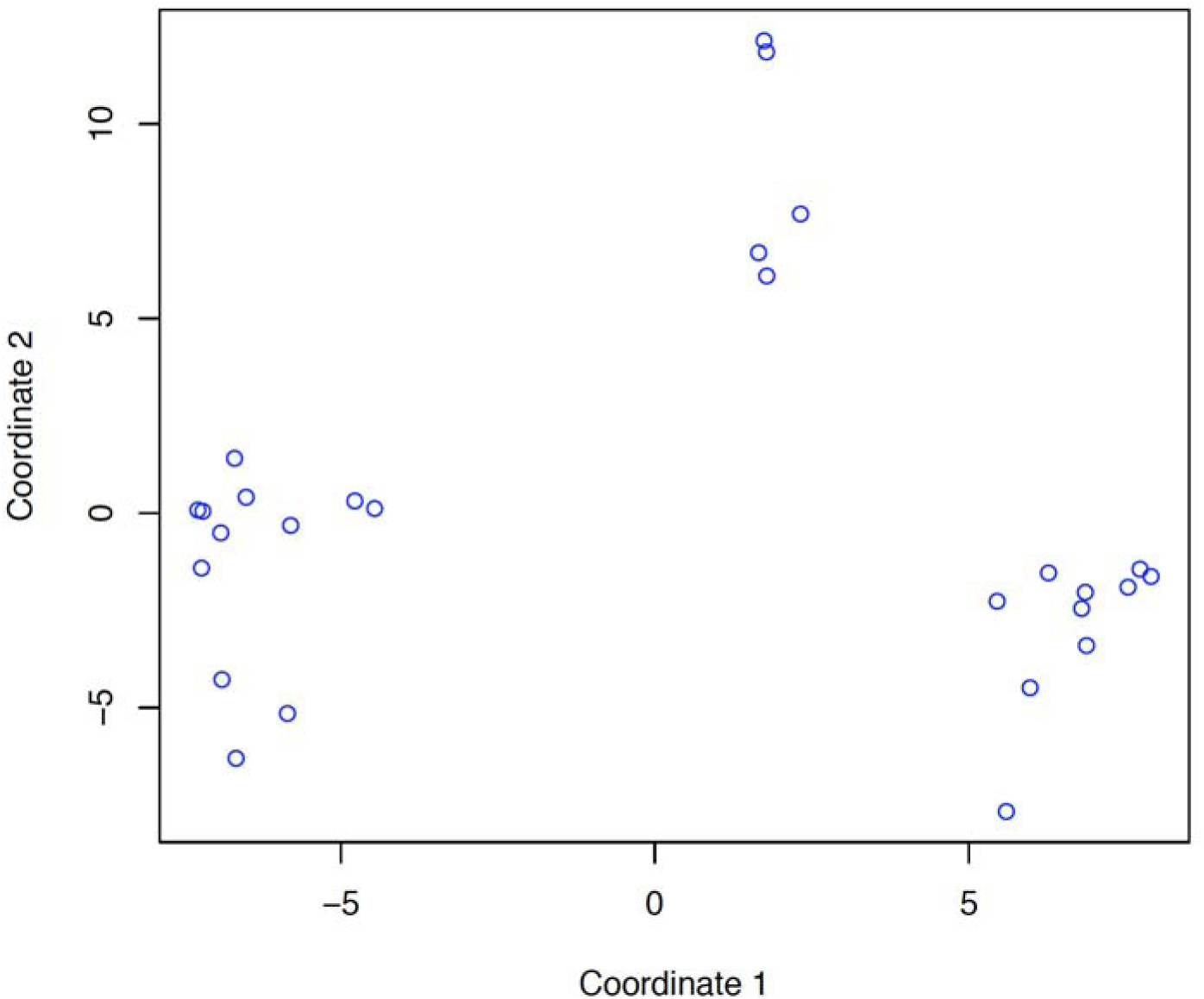
Multidimensional scaling plot of multilocus genotypes derived from SNPs in the *Porites* spp. specimens. Each point represents a specimen. Distances between points represent Euclidean distances projected in two dimensions.

**Fig. 4.**
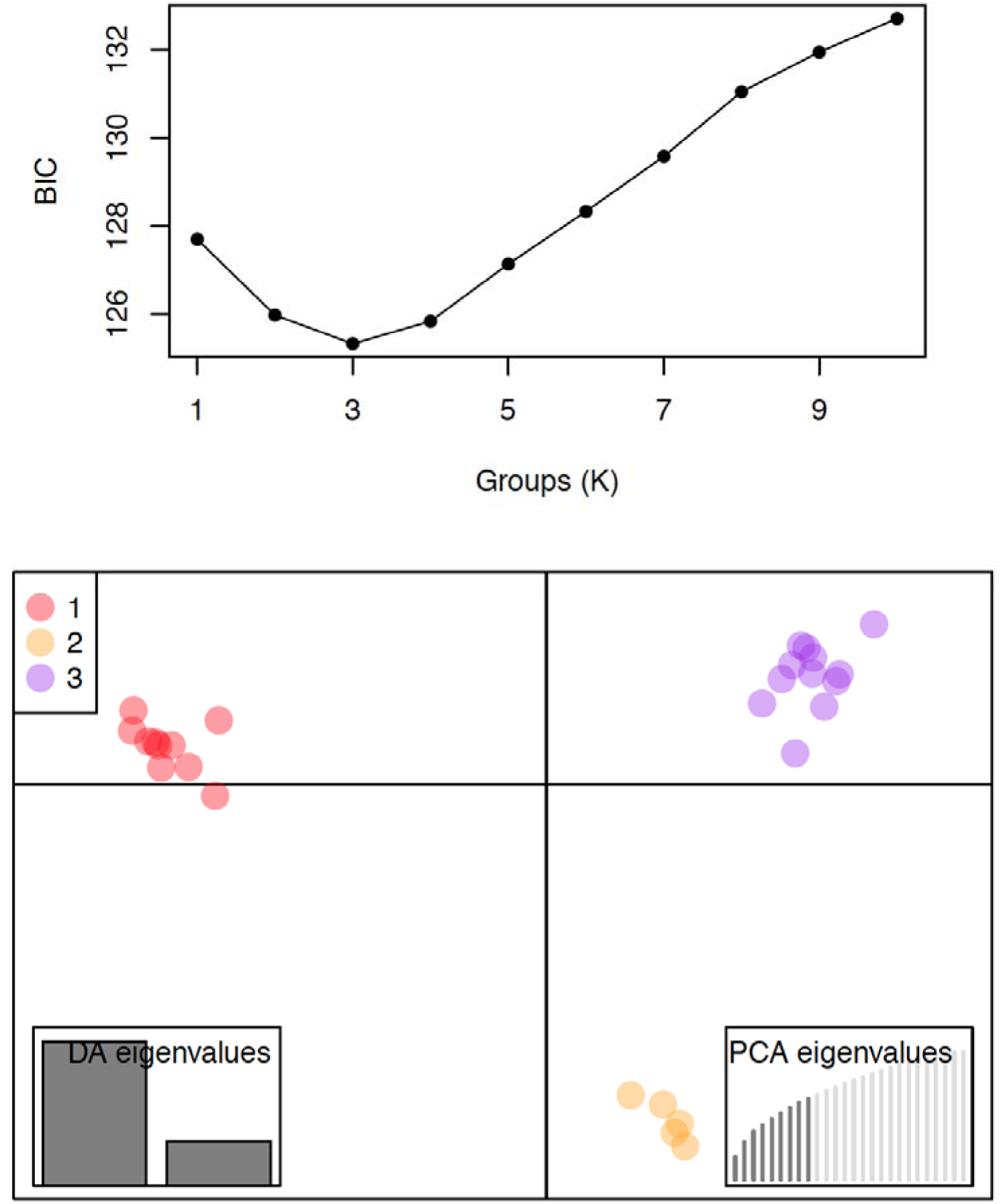
DAPC analysis of SNP data. (Top) Results of *find.clusters* analysis on SNP data identifying optimal number of groups. K = 3 groups was determined to be optimal based on the lowest value of the Bayesian information criterion (BIC). (Bottom) Results of DAPC analysis on SNP data using the three optimal groups defined by *find.clusters*. Left inset shows DA eigenvalues illustrating the relative weight of the two DA axes (x = DA 1, y = DA 2). Right inset shows the relative amount of the variance explained by the 9 principal components used in the DAPC.

We evaluated potential factors associated with genetic structure using multiple regression. For the genetic variable, we used the first discriminant axis from the DAPC analysis of SNPs. This was regressed against collection depth, symbiont type, habitat, and branch diameter. The model explained 61% of the variation in the SNP variable. The function *calc.relimp* in the R package *relaimpo* was used to estimate the relative importance of each model component. The relative importance of depth, symbiont type, habitat, and branch diameter in the model was 0.5%, 2%, 31%, and 66.5%, respectively. We further evaluated branch diameter with a one-way ANOVA, followed by pairwise t-tests with Bonferroni adjustment. Group 1 branch diameter was significantly different from both groups 2 and 3 (*p* <0.001), while groups 2 and 3 were not significantly different from each other (*p* = 0.724) (Fig. 5).

**Fig. 5.**
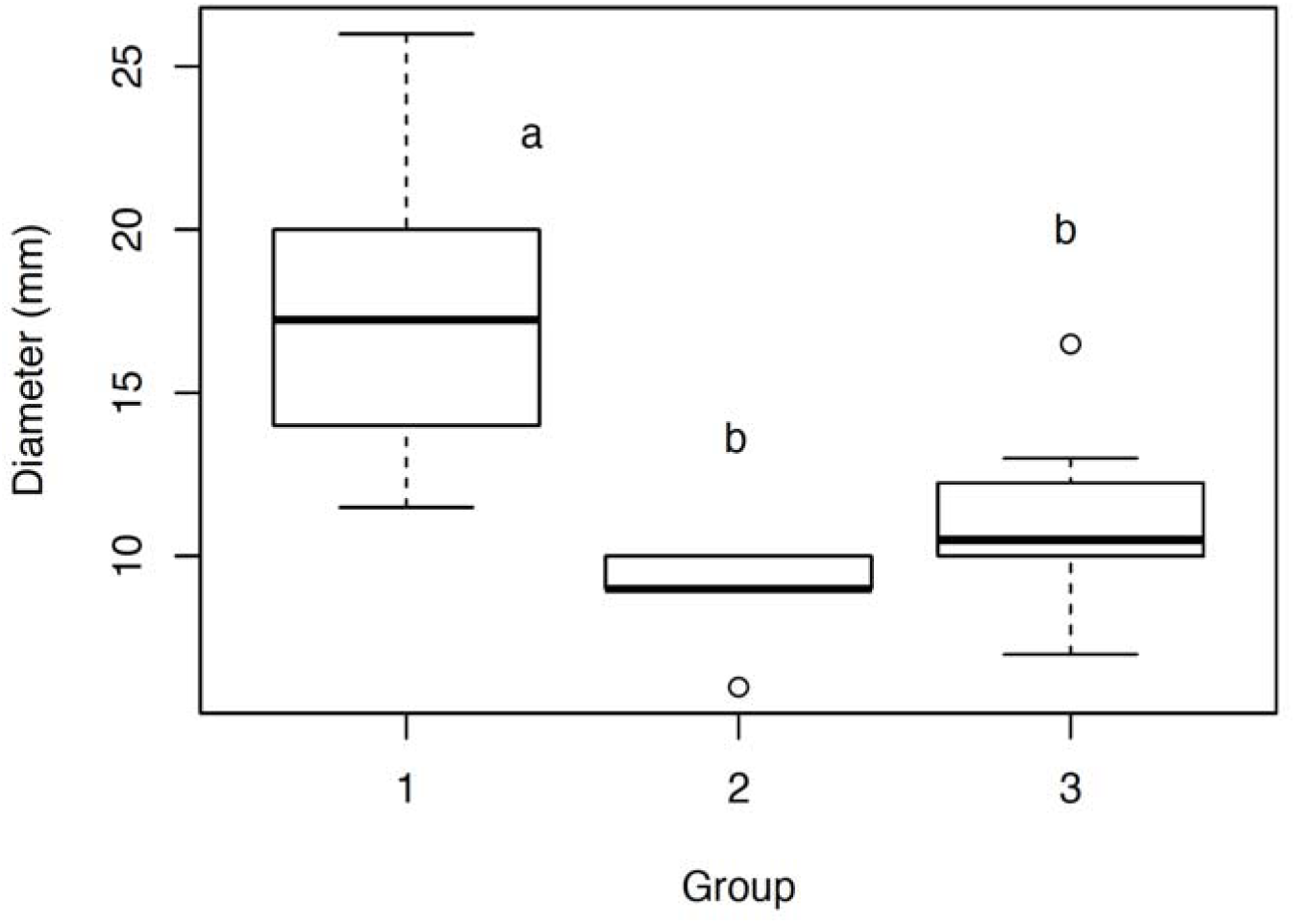
Comparison of branch diameter in the three *Porites* groups identified by the DAPC analysis in Fig. 4. Boxes show the median (black horizontal bars) plus the 75th percentile and minus the 25th percentile; whiskers show these percentiles plus or minus 1.5 times the interquartile range. Extreme values are shown as points beyond these ranges. Different letters denote groups determined to be significantly different in pairwise t-tests.

### Epigenetic patterns

Among the 1,712 CpGs, a range of 314-360 were methylated per sample, yielding a mean methylation level of 19.5% (SD 0.008%). However, there were two outliers with high levels of methylation. This was due largely to low read count loci being categorized as methylated. A more stringent minimum read count threshold of 10 reads per locus was therefore applied to the ddRADseq dataset and this resulted in 1,368 CpGs with a range of 238-263 methylated CpGs per sample, corresponding to a methylation level of 18.3% (SD 0.005%). Of the 1,368 CpGs, 208 were differentially methylated, 131 were constitutively methylated across all samples, and 1029 were constitutively non-methylated across all samples.

Methylation patterns in the MDS analysis were less clear than SNP patterns, with no clear grouping and most of the variation spread across much of the horizontal axis (Fig. 6). K-means clustering of CpG methylation indicated the strongest support for a single group (not shown), so DAPC was not appropriate. Among the 208 differentially methylated CpGs, most were either methylated or unmethylated across a majority of samples (Fig. 7). However, when visualized within the context of the groups identified in the SNP analysis, individuals in group 1 appeared to cluster adjacent to each other (far left color swatch, Fig. 7). These also tended to be samples with thicker branch diameter (bar plot, Fig. 7).

**Fig. 6.**
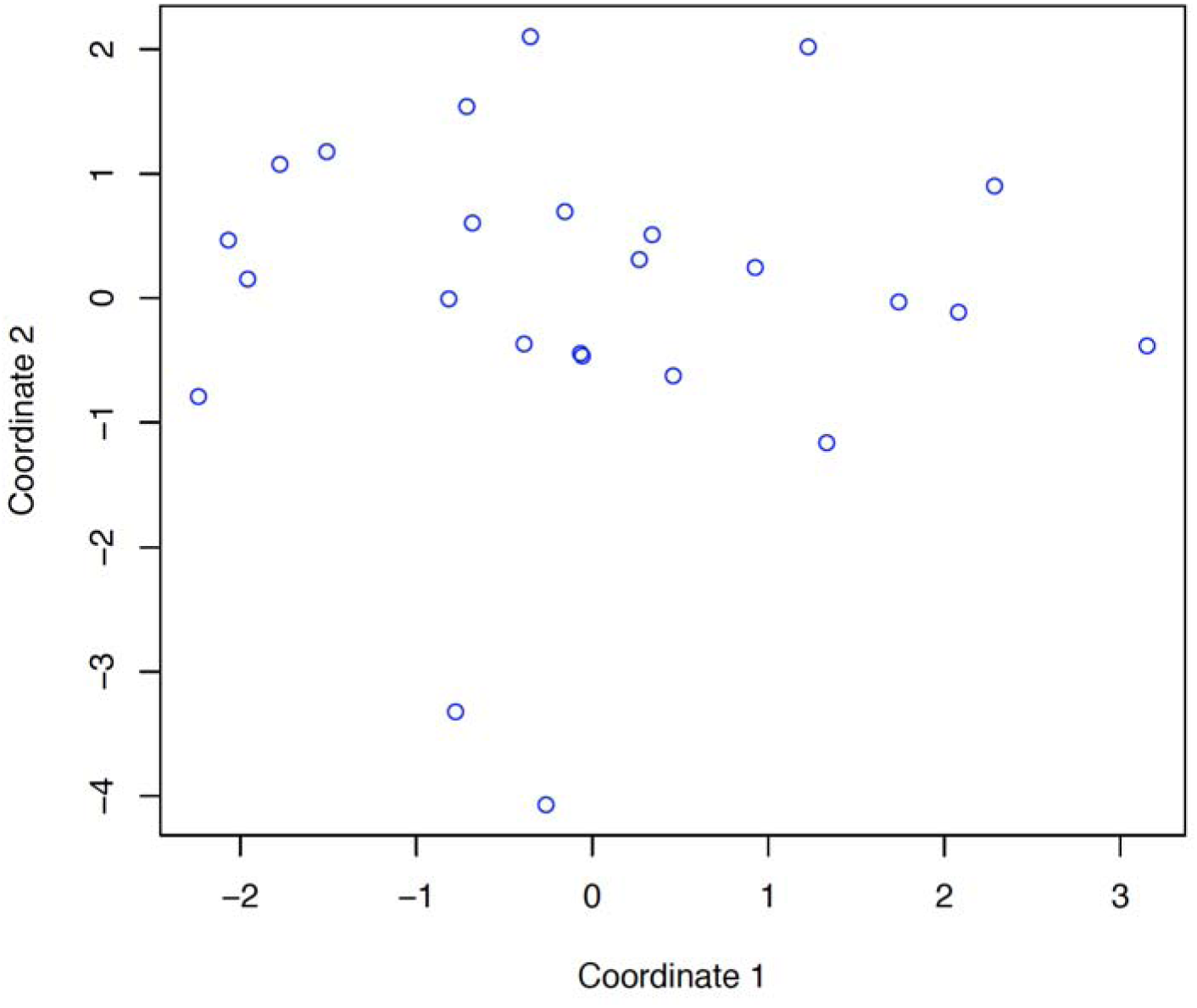
Multidimensional scaling plot of multilocus epi-genotypes in the *Porites* spp. specimens. Each point represents a specimen. Distances between points represent Euclidean distances projected in two dimensions.

**Fig. 7.**
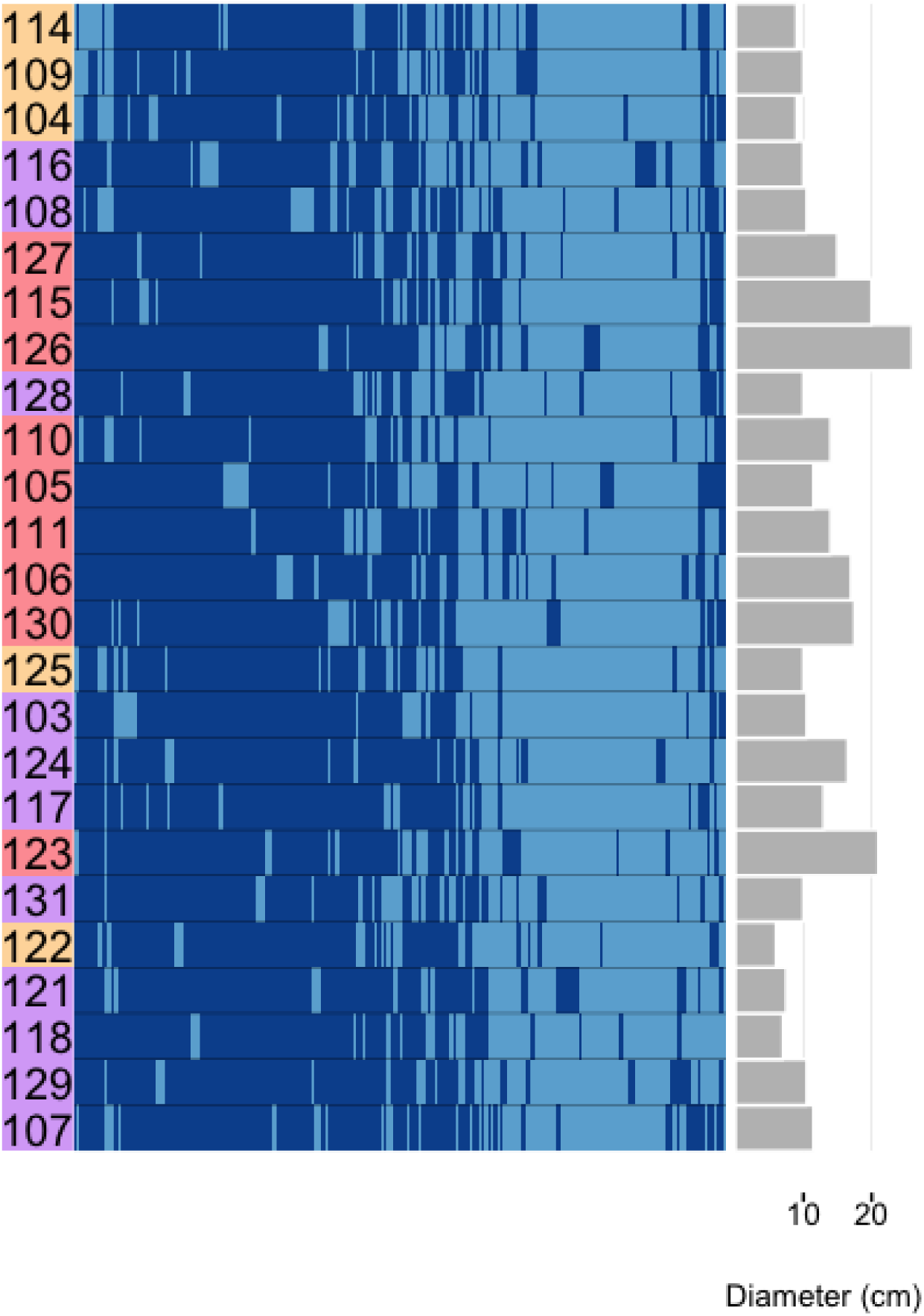
Heatmap of differentially methylated CpGs. Methylated CpGs are shown in dark blue. Hierarchical clustering was used for ordering of both rows (samples) and columns (CpGs). On the left hand side of the plot, samples are colored by the three SNP groups from the DAPC in Fig. 4. On the right side, a bar plot of the branch diameter of each sample is shown.

### Linkages between genetic and epigenetic variation

To evaluate potential associations between genetic and epigenetic variation, we asked whether SNPs with high contributions to the DAPC analysis were linked to differentially methylated CpGs (i.e., from the same RAD loci). Unique locus IDs generated by *ipyrad* were used to merge differentially methylated CpG data with the SNP contribution scores from the DAPC. There was no significant difference between contribution scores associated with differentially methylated loci and scores from a random sample of the data (Kolmogorov-Smirnov test; *p* = 0.365; Fig. 8).

**Fig. 8.**
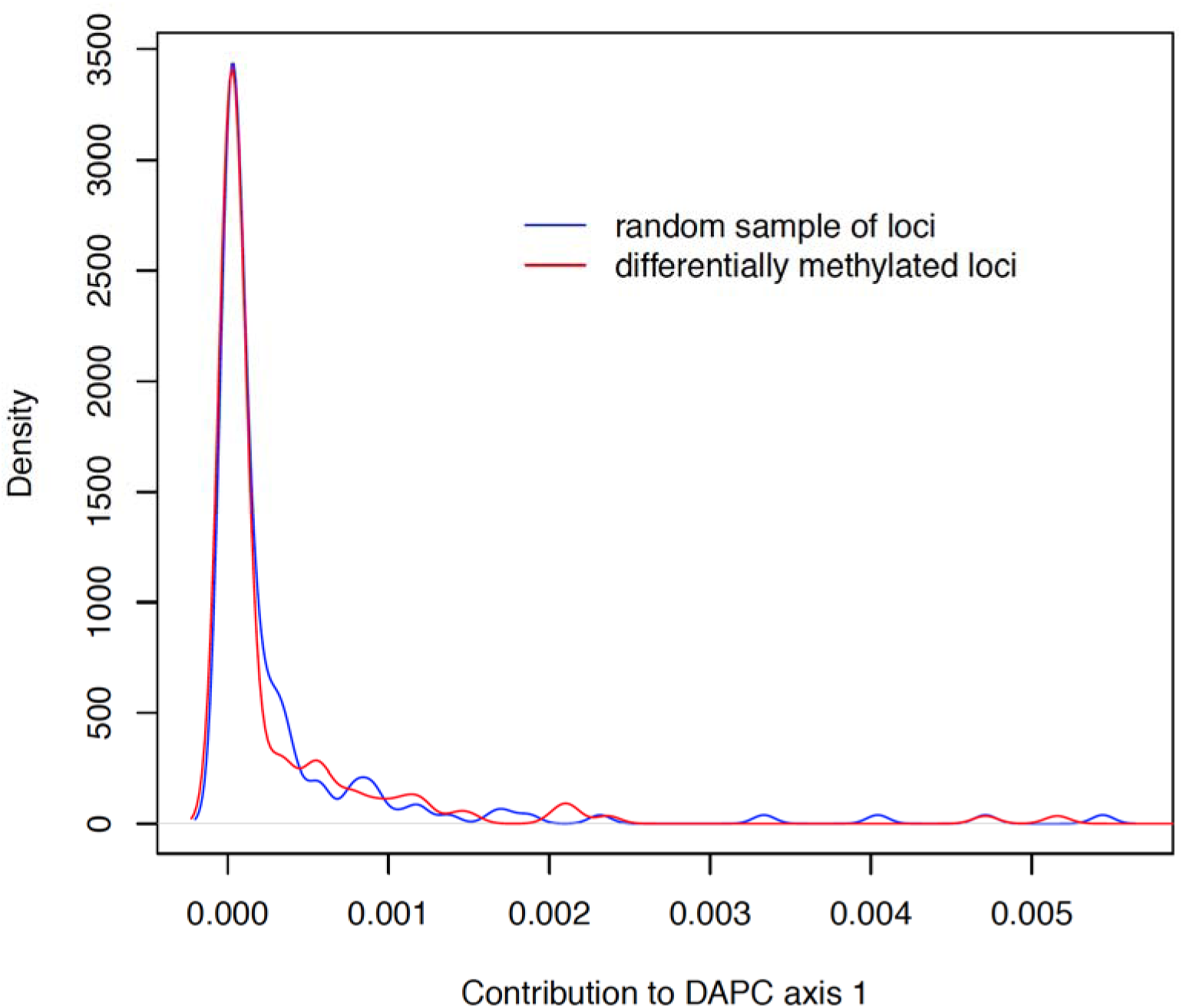
Distribution of SNP contribution scores to discriminant axis one of the DAPC model in Fig. 3. The red line shows scores of SNPs linked to differentially methylated cut sites on the same loci, while the blue line shows scores from a random sample of the data.

We also compared genetic and epigenetic variation by calculating pairwise genetic and epigenetic distances, again using the *dist.gene* function in the R package *ape*. This was the same statistic generated in Fig. 2 for the SNP data, measuring the proportion of pairwise mismatches in multilocus SNPs and CpG methylation status. The majority of pairwise comparisons did not exhibit a clear association between genetic and epigenetic distance (Fig. 9). However, the two pairs of genetic outliers identified earlier with very low genetic distances (109 & 114, 127 & 115) also had the lowest epigenetic distances. A linear regression of these data was significant (*p* < 0.001).

**Fig. 9.**
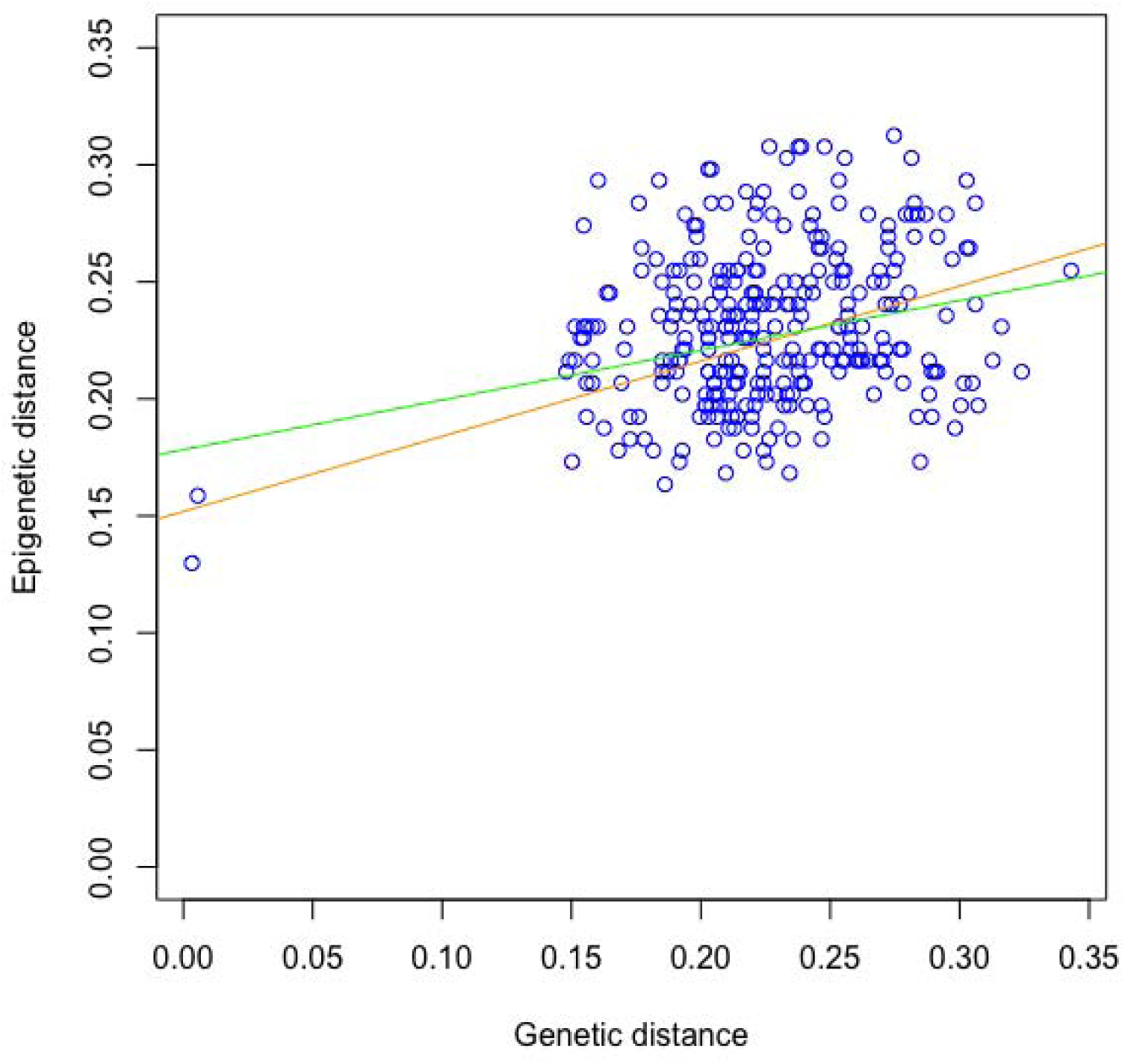
Association between pairwise genetic distance and pairwise epigenetic distance. Each point represents a unique pairwise comparison between two samples among the 25 samples that had both genetic and epigenetic data (300 unique pairwise comparisons total). For a given pair, genetic and epigenetic distances represent the proportion of mismatches between multilocus genotypes (SNPs) and multilocus epi-genotypes (CpG methylation status), respectively. The two pairs of outliers are samples 109/114 and 127/115; these possible clones showed the lowest levels of both genetic and epigenetic differentiation. Linear regressions of all the data (orange line, *p* < 0.001) and exclusive of the outliers (green line, *p* = 0.002) are shown.

## Discussion

Our sample of branching *Porites* spp. from Belize exhibited clear genetic differentiation, supporting the idea that these corals comprise three separate and fairly well-defined groups. While this result is in general agreement with older studies based on allozymes (Weil 1992; Budd *et al.* 1994), it contrasts with the most recent study based on 11 genetic markers (Prada *et al.* 2014). The inferences we were able to make likely reflect the large number of markers we sampled. Reduced representation genome sampling methods like RADseq have shown great promise in resolving phylogenetic relationships, especially in recalcitrant taxa like cnidarians for which traditional mitochondrial and nuclear markers have shown limited success (Pante *et al.* 2015; Combosch & Vollmer 2015; Herrera & Shank 2016; Rosser *et al.* 2017). Despite the number of markers we sampled, however, our study lacks sufficient geographic breadth to draw strong phylogenetic conclusions. Indeed, all evaluations of branching *Porites* spp. to date, including ours, have been limited by geographic scope, insufficient colony sample size, or low genome coverage. A thorough reexamination of branching *Porites* spp. phylogeny across the Caribbean / Western Atlantic region may be warranted, assessing both genome-wide variation and multivariate morphological traits. Ideally, such a study would incorporate randomized collections in different habitats to permit assessment of potential habitat selection.

Differences in branch diameter, particularly between group 1 and the other two groups, suggest that the genetic differentiation we observed is associated with colony-level morphological variation. Variation in branch thickness is the primary diagnostic feature used for *Porites* spp. identification in the field, and is likely what historically prompted delimitation of species groups. Corallite-level variation in morphology is also well documented, and most studies using these methods have identified distinct groups based on multivariate corallite characters (Weil 1992; Budd *et al.* 1994; Jameson 1997). Two of these studies found evidence supporting the three recognized species (Weil 1992; Jameson 1997), while one found up to five morphospecies among specimens collected from three regions (Budd *et al.* 1994), and another identified continuous variation without any clear breaks (Brakel 1977). However, despite the continuous variation observed by Brakel (1977), he judged the variation to be largely genetic (fixed) based on the observation that 1) corals from very different environments can have very similar morphologies, and 2) corals from very similar environments can have very different morphologies. This concurs with our analysis suggesting that morphotype variation in *Porites* spp. has a genetic basis. On the other hand, we also observed evidence for phenotypic plasticity, such as in the case of three specimens collected from nearby mangrove prop roots exhibiting similar branch morphologies; one of these colonies (122) was assigned to group 2 while the others (118, 121) were assigned to group 3.

Based on the lack of consensus among previous studies concerning the taxonomic status of branching *Porites* spp., and hypotheses that phenotypic plasticity might underlie this incongruence, we did not anticipate finding strong genetic differentiation and instead we hypothesized that epigenetic patterns might prove more informative. Instead, patterns of DNA methylation were less conclusive than genetic patterns. Levels of methylation did not vary greatly among samples, and even among differentially methylated CpGs, most were either methylated or unmethylated in a majority of samples. Similarly, little variation in methylation was observed in a recent study of threespine sticklebacks, with only 737 differentially methylated CpG sites identified out of 1,445,567 sites examined across eight individuals (Smith *et al.* 2015). However, these differentially methylated loci were associated with alternative phenotypes (Smith *et al.* 2015). The only phenotypic pattern to emerge from our data was a suggestion that individuals in group 1, which also tended to be more thickly branched, were generally grouped close together. Thus, there is an intriguing possibility that epigenetic variation is coupled with genetic variation in these individuals.

Genetic-epigenetic coupling is further suggested by the positive relationship between pairwise genetic and epigenetic distance. This relationship implies a potential heritable component to the methylation patterns, although it was strongest when including two pairs of outliers. The physical distances separating these pairs suggests that they are unlikely to be clones resulting from fragmentation, but possibly clones from asexually produced larvae (Harrison 2011). The interpretation that these pairs are clones is more likely, because even though some branching *Porites* spp. colonies are hermaphrodites capable of self-fertilization (Schlöder & Guzman 2008), Mendelian segregation leads to an extremely low probability of identical parental and offspring genotypes (Stoddart 1983). If they are indeed clones, the similar methylation patterns in these individuals would reflect inheritance via mitosis rather than sexual recombination through the germline. However, regardless of whether methylation profiles were transmitted sexually or asexually in the two pairs of outliers, it is notable that while genetic distance was near zero, epigenetic distance was relatively high. This could reflect divergence of the methylome due to environmental effects. Methylation levels in corals have recently been shown to be at least partially under environmental influence over short time scales (Putnam *et al.* 2016). Clearly, the methylation patterns we observed in *Porites* spp. exhibit a high degree of noise, and this noise may reflect the diverse habitat conditions and environmental histories experienced by the corals analyzed here. Environmental influence is also suggested by the case of the three corals collected from adjacent mangrove prop roots (118, 121, 122); while these corals came from two different genetic groups, they were clustered closely to each other epigenetically.

Beyond simply covarying with genetic variation, some studies have reported epigenetic variation potentially driving genetic variation (Skinner *et al.* 2015; Smith *et al.* 2016). In our analysis, differentially methylated CpGs were not more likely to be linked to SNPs with strong contributions to genetic structuring. However, it is important to keep in mind that SNPs and differentially methylated CpG sites could be separated by up to ∼500 bp on the RAD loci we generated. Moreover, given the read lengths and profiling techniques, the number of CpG sites we were able to sample per locus with EpiRADseq was a fraction of the number of nucleotides we were able to assess for SNPs. For example, for a given paired-end locus, nearly 200 nucleotides were assayed for potential SNPs whereas only a single CpG site was assayed for methylation. Nonetheless, if we consider the EpiRADseq method a random sample of CpGs, the 18% methylation we observed is within the range expected for invertebrates (Zemach *et al.* 2010; Sarda *et al.* 2012; Dimond & Roberts 2016).

Reduced representation genome sequencing techniques are currently very popular and are enhancing our ability to probe molecular processes, but they are not without error. Our analysis of ddRADseq and EpiRADseq libraries in tandem provided robust control for error in both the SNP and methylation datasets. Technical replicates have been advocated as a means to assess genotyping error, and by assessing genotypes of the same individuals from the two libraries we confirmed that this error was within the range documented elsewhere (Mastretta-Yanes *et al.* 2015; Recknagel *et al.* 2015). For the methylation analysis, comparing EpiRADseq libaries to ddRADseq libraries was a key factor in controlling for library composition effects, reducing the likelihood of false positives. This paired library approach appears to be a strong alternative to the unique molecular identifier approach used by Schield *et al.* (2016).

In addition to genetic data, prevailing symbiont populations could provide an additional means to evaluate whether branching *Porites* spp. exhibit species-level differentiation. Species-specific associations between host corals and *Symbiodinium* are common in the Caribbean, particularly among brooding species such as *Porites* (Finney et al. 2010; Bongaerts et al. 2015). Furthermore, branching *Porites* spp. appear to associate with host-specialist symbionts that are not common in other hosts (Finney *et al.* 2010; Bongaerts *et al.* 2015). Branching *Porites* spp. also tend to host distinct symbionts from their congener *P. astreoides* (Finney et al. 2010; Bongaerts et al. 2015). Although cp23S is not a fine-scale genetic marker, the majority of corals we analyzed hosted the same *Symbiodinium* clade C phylotype, while three individuals in group 3 and one in group 2 hosted the clade A phylotype (Table 1). While host-symbiont pairings can be regionally specific (LaJeunesse 2002; Finney *et al.* 2010), if branching *Porites* spp. have the same symbiont profile on a given reef regardless of potential differences in host genetics or morphology, this could be an argument against considering them separate species. A comprehensive reexamination of branching *Porites* spp. would be wise to include an assessment of *Symbiodinium* communities.

## Conclusion

Contrary to our expectations, branching *Porites* spp. morphotype variation was better explained by genetic patterns than epigenetic patterns. This analysis benefited from the resolution afforded by genome-wide sequencing, and may justify a more thorough analysis of branching *Porites* spp. phylogeny throughout the tropical Western Atlantic. Although patterns of DNA methylation were not as conclusive as genetic patterns, there was some evidence of covariation between genetic and epigenetic variation. This possibility, as well as potential environmental influence on methylation in corals, will require further study. Given the increasingly powerful molecular biology tools available for work in environmental epigenomics, stronger inferences about the extent, variability, and potential functions of epigenetic processes in corals are only a matter of time.

## Acknowledgements

We thank the Smithsonian Institution’s Caribbean Coral Reef Ecosystems Program for field support, and the Belize Fisheries Department for specimen export permitting. RADseq training and materials were generously provided by Adam Leaché and Kevin Epperly. Daniel Thornhill and Terra Hiebert provided advice on symbiont genotyping. Sam White, Hollie Putnam, Katherine Silliman, and Megan Hintz provided helpful comments that improved the manuscript. This study was supported by the Hall Conservation Genetics Research Award (UW-CoEnv), the ARCS Foundation Seattle Chapter, the John E. Halver Fellowship (UW-SAFS), and National Science Foundation Award OCE-1559940.

## Data Accessibility

DNA sequences: GenBank accession numbers KY649212-KY649238; SRA accession numbers SAMN06566335-SAMN06566364.
Repository detailing analysis methods: https://github.com/jldimond/Branching-Porites

## Author Contributions

J.D. conceived and designed the study, collected the specimens, prepared RAD libraries, analyzed the data, and drafted the manuscript. S.G. processed the specimens, extracted DNA, and performed the symbiont genotyping. S.R. contributed reagents, equipment and analysis tools.

